# Metatranscriptomic Identification of Co- and Incidental Infections in Marine Turtles Reveals Complex Disease Aetiologies

**DOI:** 10.64898/2026.10.02.753670

**Authors:** Lauren Lim, Bethan Lang, Heather Fenton, Jane Hall, Karrie Rose, Edward C. Holmes

## Abstract

Traditional veterinary pathology investigations on wild animals generally conclude a primary cause of death based on a combination of observed clinical signs, gross and microscopic lesions, microbial culture and PCR. However, these methods may fail to capture the broader microbial community of the host. We employed metatranscriptomic sequencing to characterise the potential pathobiome of necropsied marine turtles, comparing these findings with corresponding veterinary pathology reports. Tissue samples were collected from 76 individuals spanning three species of marine turtles – the green turtle (*Chelonia mydas*), the hawksbill turtle (*Eretmochelys imbricata*), and the loggerhead turtle (*Caretta caretta*) – across the coastline of New South Wales, Australia. Reported causes of mortality ranged from infectious disease to trauma. We identified a diverse range of parasites within the genera *Neospirochis*, *Hapalotrema*, *Paraorgymatobothrium*, *Eimeria* and *Callitetrarhynchus*, and bacteria including the genera *Aeromonas*, *Clostridium*, and *Vibrio*. Novel viruses of the families *Astroviridae*, *Caliciviridae*, *Hepaciviridae*, *Hepeviridae*, *Nyamivirdae*, *Orthomyxoviridae*, *Picobirnaviridae* and *Rhabdoviridae* were also identified. In cases where infectious disease was noted in the pathology report, we frequently observed high pathogen read counts consistent with the listed primary diagnoses. However, we also detected a range of incidental infections involving eukaryotic parasites, bacteria, and viruses. The diversity of infectious agents detected in marine turtles suggests that the observed disease outcomes are unlikely to be driven by a single causative agent, but rather by complex interactions between host health, the pathobiome, and environmental stressors. We discuss the potential implications for turtle health and conservation.

## Introduction

Marine turtles (*Cheloniidae*) are found globally in every ocean and capable of travelling thousands of kilometres in a single migration season, limited only by the width of ocean basins and polar temperatures (Hays & Scott, 2013). As a result, marine turtles exhibit some of the broadest geographic ranges relative to body size of any species. Marine turtles are also a keystone species within marine ecosystems, maintaining sea grass meadows and regulating populations of jellyfish and sponges in coral reefs (Hendrix & Pérez-Espona, 2024). Their wide occupation of marine environments and ecological significance highlights the importance of their conservation and continued surveillance for health and disease. An assessment of the role played by marine turtles as hosts, vectors, and reservoirs for pathogens can also provide crucial insights into the dynamics of infection and transmission within marine ecosystems, in turn informing conservation strategies.

Disease surveillance efforts in marine turtle populations are limited by remote capture sites, postmortem changes, and the difficulty of investigating disease in controlled experimental studies due to complex husbandry requirements and the endangered status of many species. Despite this, marine turtles are known to suffer from a range of parasitic, fungal, bacterial and viral pathologies (Mashkour et al., 2020). Bacterial infections tend to be associated with opportunistic pathogens often reported as commensal microbiota in diet related species such as fish and crustaceans. Commonly reported bacterial taxa implicated in opportunistic or pathogenic infections include *Vibrio* spp., *Pseudomonas* spp., *Enterococcus* spp., *Aeromonas* spp., *Escherichia coli*, and *Providencia* spp. (Fichi et al., 2016; Orós et al., 2005; Zavala-Norzagaray et al., 2015). Similarly, fungal pathogens of marine turtles are typically opportunistic, with *Fusarium* species most frequently reported in nests, eggs and adult abscesses (Gleason et al., 2020; Sarmiento-Ramírez et al., 2014; Williams et al., 2012). Marine turtles are also known to harbour a diversity of eukaryotic parasites such as the apicomplexan *Eimeria* (Chapman et al., 2016), and platyhelminths such as spirorchiid trematodes, including *Neospirorchis* spp. (Marangi et al., 2024), *Hapalotrema* spp. and various other trematodes (Chapman et al., 2019; Santoro et al., 2006), although whether they serve as the definitive hosts remains unclear. In contrast, far less is known about viral infections in marine turtles, with Chelonid alphaherpesvirus 5 (ChHV5), the agent of fibropapillomatosis, currently the only virus known to cause disease in marine turtles (Ackermann et al., 2012; Herbst, 1994). While a systemic herpes virus-like disease has been reported (Curry et al., 2000; Jacobson et al., 1986), the causative virus remains uncharacterised. Severe meningoencephalomyelitis perhaps associated with piscichuvirus infection has also been documented (Laovechprasit et al., 2023). Reports of viruses in gastrointestinal samples from marine turtles necessarily reflect those of dietary origin rather than viruses infecting the turtles themselves (Laovechprasit et al. 2026).

Pathology investigations on wild animals generally identify a primary cause of death based on a combination of observed clinical signs prior to death, gross and microscopic lesions, microbial culture and targeted PCR. While these methods provide insights, they may fail to capture the broader microbial community, including taxa that potentially influence host health and disease dynamics. Metatranscriptomic (i.e., total RNA sequencing) approaches have become increasingly valuable in wildlife health surveillance by offering a potentially unbiased view of microbial activity, in turn greatly expanding our knowledge of the diversity of bacteria, fungi, viruses and parasites within hosts (Harvey et al., 2026; Nakagawa et al., 2023; Shakya et al., 2019).

Interactions between marine turtles and humans predominantly occur in coastal communities adjacent to nesting beaches, where these animals contribute economic value through ecotourism and play culturally significant roles. Within such communities, the consumption of turtle products has also resulted in cases of chelonitoxism (i.e., food poisoning), for which the aetiological agent remains unknown (Semmouri et al., 2024). Together, these human-animal interactions demonstrate the broader One Health context in which marine turtle health and human well-being intersect (Pitt & Gunn, 2024), further emphasising the need for continued wildlife surveillance.

Herein, we used metatranscriptomic sequencing to characterise the pathobiome of 76 marine turtles spanning three species: the green turtle (*Chelonia mydas*), the hawksbill turtle (*Eretmochelys imbricata*), and the loggerhead turtle (*Caretta caretta*), sampled from various locations along the east coast of New South Wales, Australia. We report a wide diversity of eukaryotic parasites, bacteria and viruses, assess differences in pathobiome composition between individuals, and integrate findings from metatranscriptomic sequencing and veterinary pathology reports to better understand the marine turtle pathobiome.

## Methods

### Animal Ethics

Marine turtle carcasses were transferred to the Australian Registry of Wildlife Health at Taronga Zoo, Sydney, Australia, for examination and sample collection in accordance with NSW National Parks and Wildlife Act 1974, section 132c, Scientific License number SL100104.

### Sample Collection

Marine turtles were collected opportunistically across the east coast of NSW, Australia (Figure 1A) between 2008 and 2021 (Supplementary Table 1). A total of 76 individual turtles were sampled, comprising: green turtle (*Chelonia mydas*; GST, n=54), hawksbill turtle (*Eretmochelys imbricata*; HBT, n=17) and loggerhead turtle (*Caretta caretta*; LHT, n=5). Broadly, case mortalities were reported due to infection (n=36; 47%), entanglement or entrapment (n=13; 17%), trauma (n=8; 10%), foreign body (n=6; 7.8%), or unknown (n=13; 17%). All animals were necropsied and tissue samples snap frozen at −80°C. Tissue availability varied between cases and included liver, lung, kidney, spleen, brain, gastric tissue and thyroid (Supplementary Table 2).

**Figure 1.**
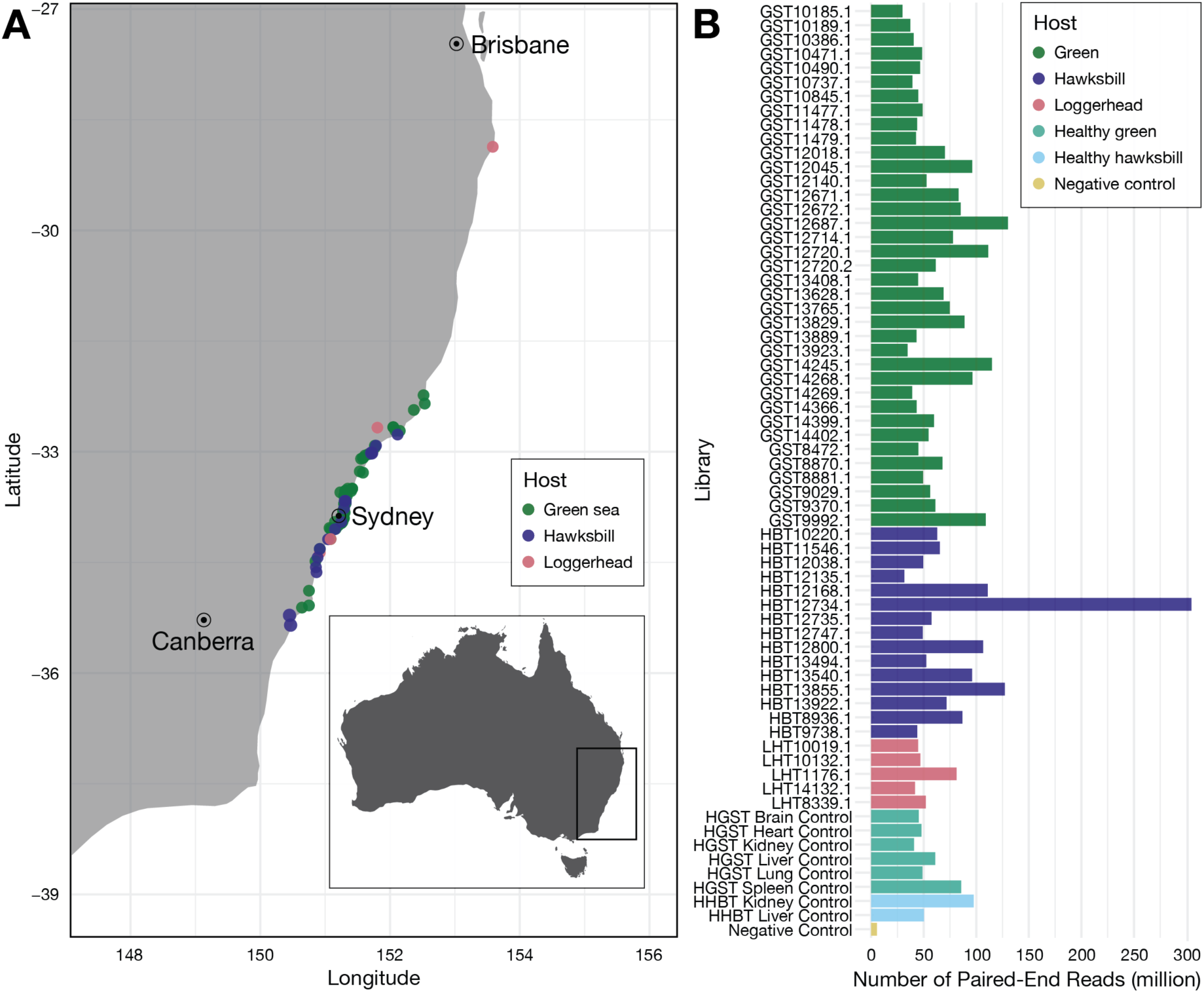
Sample collection locations and data summary of marine turtle libraries. (A) Marine turtle sampling sites across the east coast of New South Wales, Australia. (B) Total number of paired-end reads in each sequencing library. More information on individual sequencing libraries and exact sample locations is presented in Supplementary Table 1.

### Veterinary Assessment

Each turtle was subject to routine gross post-mortem examination, sample collection, and histopathological examination. Fresh portions of brain, lung, liver, skin, heart, kidney and any lesions were collected aseptically post-mortem and frozen at −80°C. A range of tissues from each animal was fixed in 10% neutral buffered formalin, processed in ethanol, embedded with paraffin, sectioned, stained with hematoxylin and eosin, and mounted with a cover slip for examination by light microscopy. In some instances, various special stains were applied to additional tissue sections to further characterise organisms seen microscopically. Additional diagnostic techniques using fresh tissues such as aerobic, anaerobic and fungal culture, and organism identification using mass spectrometry techniques, were undertaken to identify pathogens observed grossly or histologically.

### RNA Extraction

RNA was extracted from the turtle samples using the QIAGEN RNeasy Plus Mini kit (QIAGEN, Hilden, Germany) according to the manufacturer’s instructions. Tissue samples were homogenised using the TissueRuptor (QIAGEN, Hilden, Germany). A negative control was established by running buffer through clean columns to ensure no contamination occurred during the extraction process. Designed healthy control animals comprised individuals whose deaths were reported to be by acute physical trauma with no evidence of infection or disease identified upon gross and microscopic post-mortem examination. Of note, this healthy control group excluded individuals that died of trauma with the carapace or body cavity broken into (e.g., boat strike) due to possible environmental infiltration which could have biased results. This assigned 57 turtles as individual cases of varying pathology, while 19 individuals were assigned to the healthy control group and pooled by species and tissue type to form a set of 8 healthy control libraries.

RNA was quantified using the Qubit fluorometer (ThermoFisher Scientific, Massachusetts, USA) and pooled samples were further tested for DNA contamination using a parallel capillary electrophoresis instrument (TapeStation Fragment Analyzer; Agilent, California, USA).

### Total RNA Sequencing

Total RNA sequencing of the extracted RNA was conducted using the NovaSeqX platform (150bp paired end; Illumina, California, USA) by the Australian Genome Research Facility (AGRF, Melbourne, Australia). Ribosomal RNA was depleted using the Illumina rRNA depletion kit, prepped with Illumina Truseq Total RNA Library Preparation Protocol. A total of 66 sequencing libraries: 57 individual turtle libraries, eight pooled healthy control libraries and a negative (“blank”) control was sequenced. The raw sequence data was trimmed of its Illumina adapters using Trimmomatic (v.0.35; Bolger et al., 2014) in the configuration “LEADING:3 TRAILING:3 MINLEN:36”. FastQC (v.1.9.; Andrews, 2010) on raw data and trimmed paired reads evaluated libraries as high quality and confirmed successful trimming. To assess the prevalence and significance of the sequencing data, pathology reports were used to supplement the findings of our analyses throughout each step of the methods.

### Classifying fungal and non-fungal eukaryotic reads, and defining the pathobiome

Taxonomic assignment and the sorting of paired trimmed reads was achieved using the Kraken2 (v.2.1.3) metagenomic taxonomic classifier (Wood et al., 2019), with the confidence threshold flag set at 0.2 and the core nucleotide or “core_nt” database (as of June 2025). Taxa with fewer than 20 reads were removed on the assumption that they likely represented misassigned reads or very low abundance noise. The Pavian program was then used to filter and navigate the large volume of metatranscriptomic data (Breitwieser & Salzberg, 2019). The Kraken2 output was also used to verify that sequencing profiles reflected a high abundance of Sauropsida reads as expected for turtles and to identify potential human contamination.

We define the ‘pathobiome’ as all infectious agents with the potential to act as pathogens, including opportunistic taxa. Eukaryotic reads were first filtered to exclude all Chordata, which primarily represented host (Sauropsida) or dietary (Actinopterygii - fish) sequences. Further refinement excluded other phyla associated with the omnivorous diet of marine turtles, specifically Arthropoda, Mollusca, Cnidaria, Porifera, Ctenophora, Echinodermata and Viridiplantae. The remaining phyla were then sorted by maximum raw read counts and those with fewer than 40 reads (Annelida, Cercosoa, Evosea) were excluded as likely environmental background noise. Due to the high diversity of fungal taxa in the libraries, the eukaryotic component of each library was divided into two profiles: (i) the mycobiome (fungi) and (ii) all remaining eukaryotes, hereafter referred to as the parasitome. Even though some phyla included in the parasitome are likely to be non-pathogenic environment-associated species, they are included in the analysis as some species variants have been reported to cause disease. This includes amoeba and members of neurotoxin producing taxa such as Bacillariophyta (diatoms) that can negatively affect marine life.

Parasite reads were extracted using the KrakenTools suite (https://github.com/jenniferlu717/KrakenTools/tree/master) and mapped to 28S rRNA sequences using BBMap (v.38.93; Bushnell, 2014) for confirmation. We limited our analysis to parasites with high read abundance and robust read coverage across reference parasite 28S sequences (Supplementary Table 3). The *ggplot2* package (v3.5.1) in R (v4.3.3) was utilised for data visualisation (Supplementary Figure 2). When read mapping failed to provide substantive evidence of parasite presence, we attempted to use MEGAHIT *de novo* assembled contigs from the virus analysis (see below) to conduct an unbiased search for 28S sequences. A custom nucleotide turtle parasite 28S database was constructed by taking representative reference sequences from NCBI using DIAMOND BLASTx. To remove false positive sequences, candidate parasite contigs were then screened against the BLAST Nucleotide database (NCBI databases from August 2025; v.2.10.1; Sayers et al., 2024).

### Bacterial reads and the presence of bacterial toxins

Because RNA-sequencing and rRNA depletion limits bacterial taxonomic resolution, Kraken-derived bacterial classifications were restricted to genus level specificity. The ambiguity of bacterial pathogenicity at the genus level in turn meant that reads mapping to obligate, facultative and opportunistic taxa were retained without discrimination for the pathobiome analysis. Hence, common laboratory contaminants present in our negative control, such as *Ralstonia* and *Escherichia* (Salter et al., 2014; Xiao et al., 2019), were retained as they are also known to have pathogenic strains. Taxonomic profiles were generated by extracting bacterial relative abundances at the genus level using Pavian. Low abundance bacterial genera (≤1% relative abundance) were removed, and the data renormalised and plotted in R (v4.3.3) using the *ggplot2* (v3.5.1), *dplyr* (v1.1.4), and *tidyr* (v1.3.1) packages to produce the final bacterial taxa profile.

To compare variation in bacterial community composition, we computed beta diversity using the *vegan* (v2.6-8), *ggplot2* (v3.5.1), and *dplyr* (v1.1.4) packages in R (v4.3.3). We calculated Bray-Curtis distance as a measure of dissimilarity between sequencing libraries which were then visualised using principal coordinate analysis (PCoA) plots. To identify libraries with markedly divergent bacterial profiles we statistically defined “outlier” libraries, representing those that had >4 median absolute deviations (MAD) from the median of the reported Bray-Curtis distances. Median distance was chosen because of its resistance to the impact of outliers. To determine whether outlier classification was sufficiently robust a sensitivity analysis was conducted across 2.5-5 MAD. This resulted in the identification of 15 libraries that had distinct bacterial compositions (see Results; Supplementary Table 4).

Libraries from septic turtles exhibiting distinct microbiome profiles were further interrogated using the Toxinome database (flags “-e 1E-20, --sensitive” Danov et al., 2023) to identify toxin-associated sequences within the assembled contigs. Putative toxin-like sequences exhibiting ≥80% amino acid identity to known toxins were manually validated using Geneious Prime (v.2025.1.) and web-based BLASTx searches (Supplementary Table 5). As our knowledge of reptilian cellular responses to bacterial toxins is limited, only sequences associated with mammalian cytopathic effects were retained, with other toxin-related hits from the Toxinome database excluded.

### Virus analysis

Trimmed reads were assembled using MEGAHIT (v.1.2.9; Li et al., 2015). The number of reads per contig was estimated using RNA-Seq by Expectation-Maximisation (RSEM-v1.3.1; Bowtie2-v2.5.4; Salmon-v1.10.3; Samtools-v1.22.1) and reported in the number of transcripts per million (TPM). Contigs were screened against the clustered Reference Viral Database (as of May 2025; Bigot et al., 2020) and RdRp-Scan (Charon et al., 2022) using DIAMOND BLASTx (v.2.16.0; flags set at “-e 1E-5 --ultra-sensitive”; Buchfink et al., 2015). Further screening was performed using DIAMOND BLASTx against the NCBI non-redundant (nr) protein database (NCBI databases from August 2025; v.2.16.0; Sayers et al., 2024) to identify and remove false-positive contigs. This screening process resulted in 256 potential viral contigs that were then organised by viral order or family and subjected to ORF prediction using Geneious (v.2025.1.). Finally, a manual web-based BLAST analysis was used to confirm virus-like contigs >699bp in length. A total of 90 viral contigs were confirmed using this method (Supplementary Table 6).

To infer phylogenetic trees of each RNA virus order or family identified we first constructed RNA-dependent RNA polymerase (RdRp) amino acid alignments using MAFFT E-INS-i algorithm (Katoh & Standley, 2013). Background reference virus sequences to provide phylogenetic context were taken from NCBI/GenBank and sequence alignments trimmed using trimAl (Capella-Gutiérrez et al., 2009). Phylogenetic trees were then estimated using the maximum likelihood (ML) method available in IQ-TREE2 (Minh et al., 2020) with the ‘ModelFinder Plus’ (-m MFP) flag used to select the best-fit model of amino acid substitution (Kalyaanamoorthy et al., 2017), 1000 ultrafast bootstrap replicates, and optimised by nearest neighbour interchange (-bnni). Phylogenies were further annotated using FigTree (v1.4.4.; Rambaut, 2018) and edited using Adobe Illustrator (2025) and the R (v4.3.3) *ggtree* package (v3.8.2).

## Results

Our metatranscriptomic analysis of 76 marine turtles generated 4,400,810,764 paired-end reads with a range of 29,865,874 – 304,621,949 read, and totalling 1.3Tb of data over 65 total RNA-sequencing libraries excluding the negative control (the latter with 5,501,434 paired-end reads; 1.65 Gb). On average, 66,678,950 paired-end reads and 20.0 Gb of data was generated for each sequencing library (Figure 1B). Kraken2 classified 54.5-95.0% reads in each library (average 73%), of which 77.9-97.52% were attributed as chordate reads (average 92%). Testudine reads comprised 22.75-82.89% of the sequencing libraries and were the most abundant animal group in every case. Neither the negative control nor any of the 65 turtle libraries contained human contamination or any other unexpected signals. Cases with a lower relative abundance of testudine reads and higher bacterial reads were associated with cases of sepsis, pneumonia and trauma to the carapace.

For most libraries, the mycobiome component comprised high fungal taxonomic diversity but low fungal read abundance (i.e., an average of 804 reads across 65 libraries). Fungal genera commonly reported in marine or environmental contexts (including *Nakseomyces*, *Fusarium*, *Heterosporis*, *Malassezia*, *Penicillium*) were not dominant in any library and only detected at low relative abundance. While some of these candidate fungal pathogens are known opportunistic pathogens of marine turtles, their detection here occurred in cases where fungal infection was not observed as the primary cause of death and hence likely reflect an incidental finding. Consequently, the mycobiome component was deemed of low importance and not included in downstream analyses.

### Eukaryotic parasites of marine turtles

After taxonomic classification and read map confirmation against 28S rRNA reference sequences we identified six parasite genera in 24 sequencing libraries (Supplementary Figure 2). Consistent with previous reports of marine turtle parasites, platyhelminth blood flukes (Spirorchiidae) of the genera *Neospirorchis* and *Hapalotrema* were most prevalent in our data – detected in 15/65 (23.1%) and 8/65 (12.3%) libraries, respectively (Table 1). *Neospirorchis* was detected in both green and hawksbill turtle libraries, while *Hapalotrema* was detected in green and loggerhead turtle libraries.

**Table 1.**
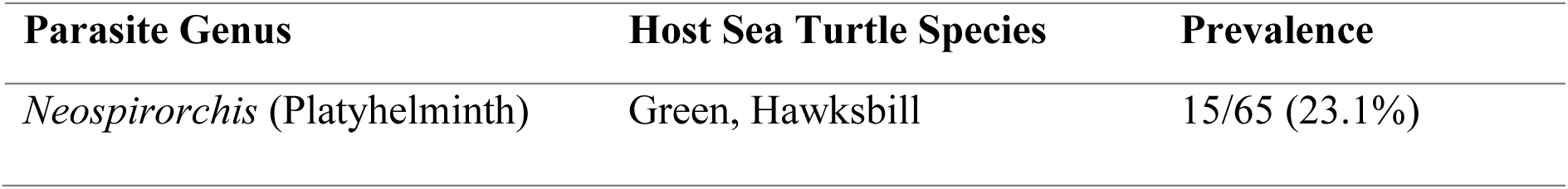
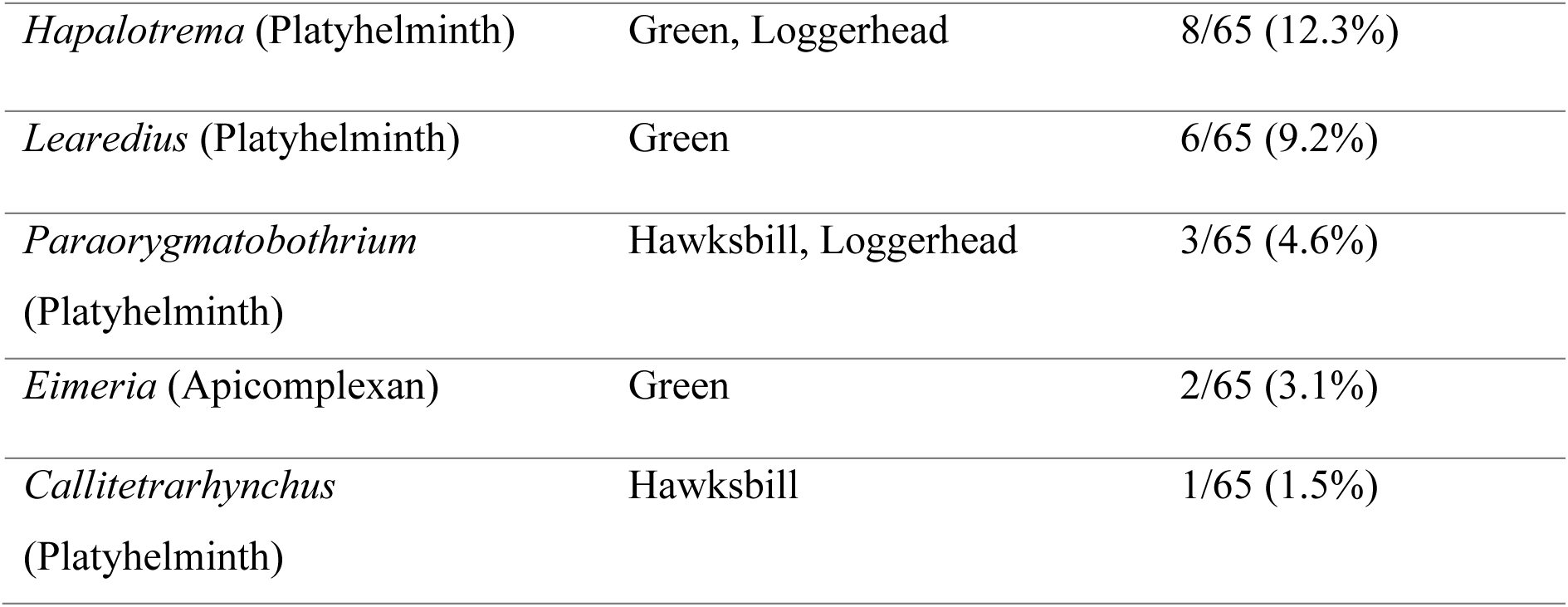
Prevalence of eukaryotic parasite-associated detections across marine turtle sequencing libraries. Prevalence in total number of libraries.

Severe infection with *Learedius* has been reported to cause blindness in green turtles of the northern Red Sea (Morick et al., 2023), and was detected in 6/65 libraries (9.2%). This likely represents an incidental finding as none of the six cases in our study were associated with turtle blindness. Lower prevalence detections included *Paraorygmatobothrium* in 3/65 libraries (4.6%), *Eimeriia* in 2/65 (3.1%) and *Callitetrarhynchus* in a single library (1/65; 1.5%). While trematode detections are consistent with known marine turtle-associated parasites, the presence of *Paraorygmatobothrium* and *Callitetrarhynchus* should be interpreted with caution as these taxa are more commonly associated with fish and may therefore reflect transient dietary or environmental sources.

Of the 24 parasite-positive libraries, parasites were recorded as the primary cause of disease or mortality in five cases due to the high parasite burden observed (Supplementary Table 3), while a further nine cases included pathological observations of parasites or parasite-associated lesions (Figure 2). In the remaining 10 parasite-positive libraries, parasites were not noted in the available pathology records and therefore more likely represent incidental findings. Together, these metagenomic data indicate that marine turtle libraries contain diverse eukaryotic parasite-associated sequences, although the detection of parasite reads alone does not confirm an aetiologic involvement in individual turtle cases.

**Figure 2.**
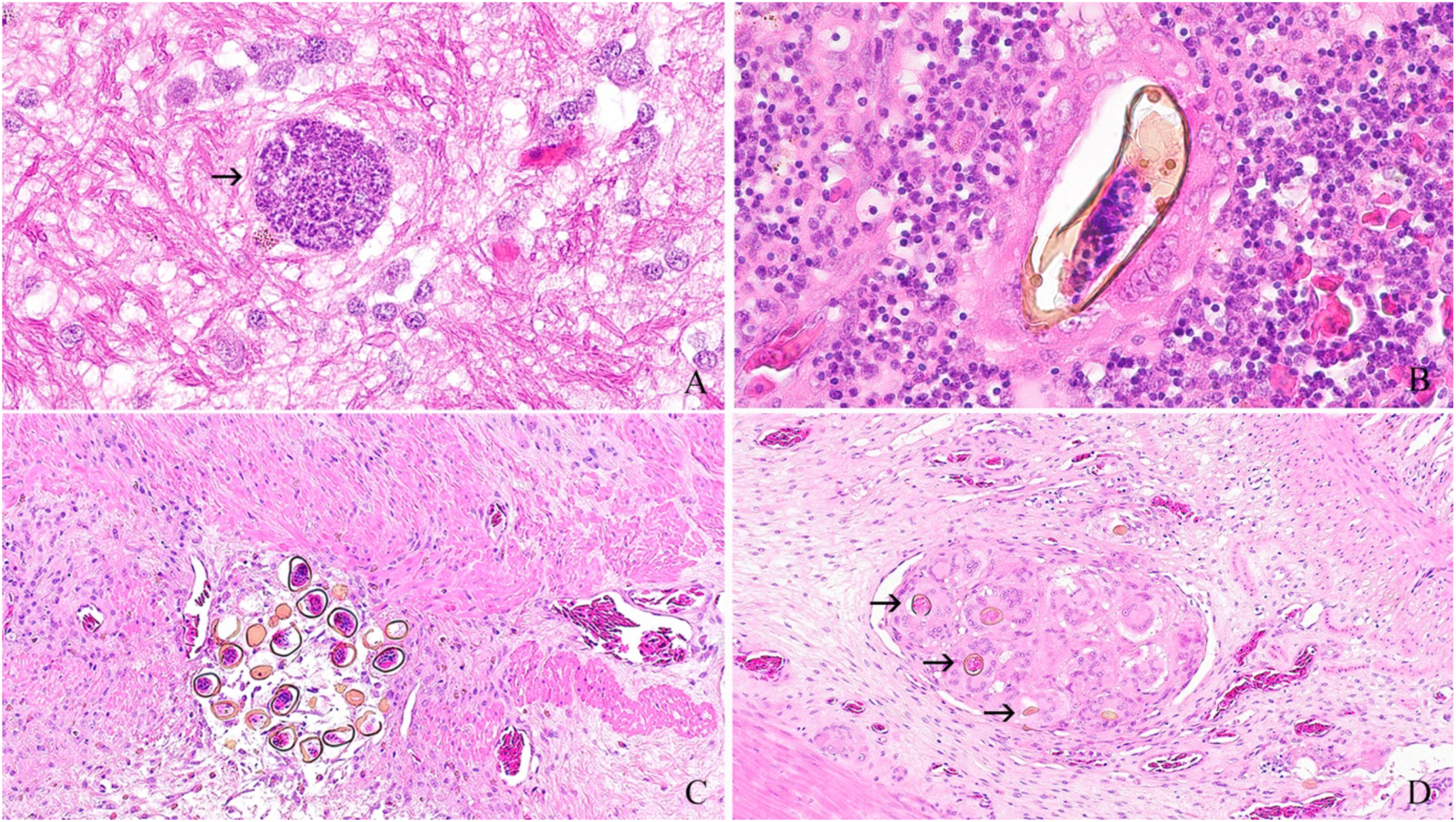
Histological findings in marine turtles. (A) Brain from a subadult green turtle (*C. mydas*) with an Eimeridae protozoal meront (arrow) surrounded by loose myelinated fibers, vacuolation of the neuropil, and small aggregates of glial cells. Hematoxylin and eosin stain at 40x objective. (B) Single spirorchid trematode egg with a thick, refractile yellow shell attributed to *Neospirorchis* sp. surrounded by a granuloma composed of multiple epithelioid macrophages and multinucleated giant cells in the spleen from a subadult green turtle. Hematoxylin and eosin stain at 40x objective. (C) Granuloma with multiple spirorchid trematode eggs surrounded by epithelioid macrophages and multinucleated giant cells within the muscularis of a section of small intestine from a subadult green turtle. Hematoxylin and eosin stain at 40x objective. (D) Granuloma with multiple spirorchid trematode eggs (arrows) surrounded by epithelioid macrophages and multinucleated giant cells within the submucosa of a section of small intestine from a subadult green turtle. Hematoxylin and eosin stain at 40x objective.

### Bacterial infections within marine turtles

Bacterial reads were the most abundant within the pathobiome, ranging between 8,531-6,237,043 and averaging 423,379 Kraken classified reads per library. Although the most abundant bacterial genus detected across all sequencing libraries was *Ralstonia* (Figure 3A), it was also detected at high relative abundance in the negative control suggesting that its presence likely results from reagent contamination as noted previously (Salter et al., 2014).

**Figure 3.**
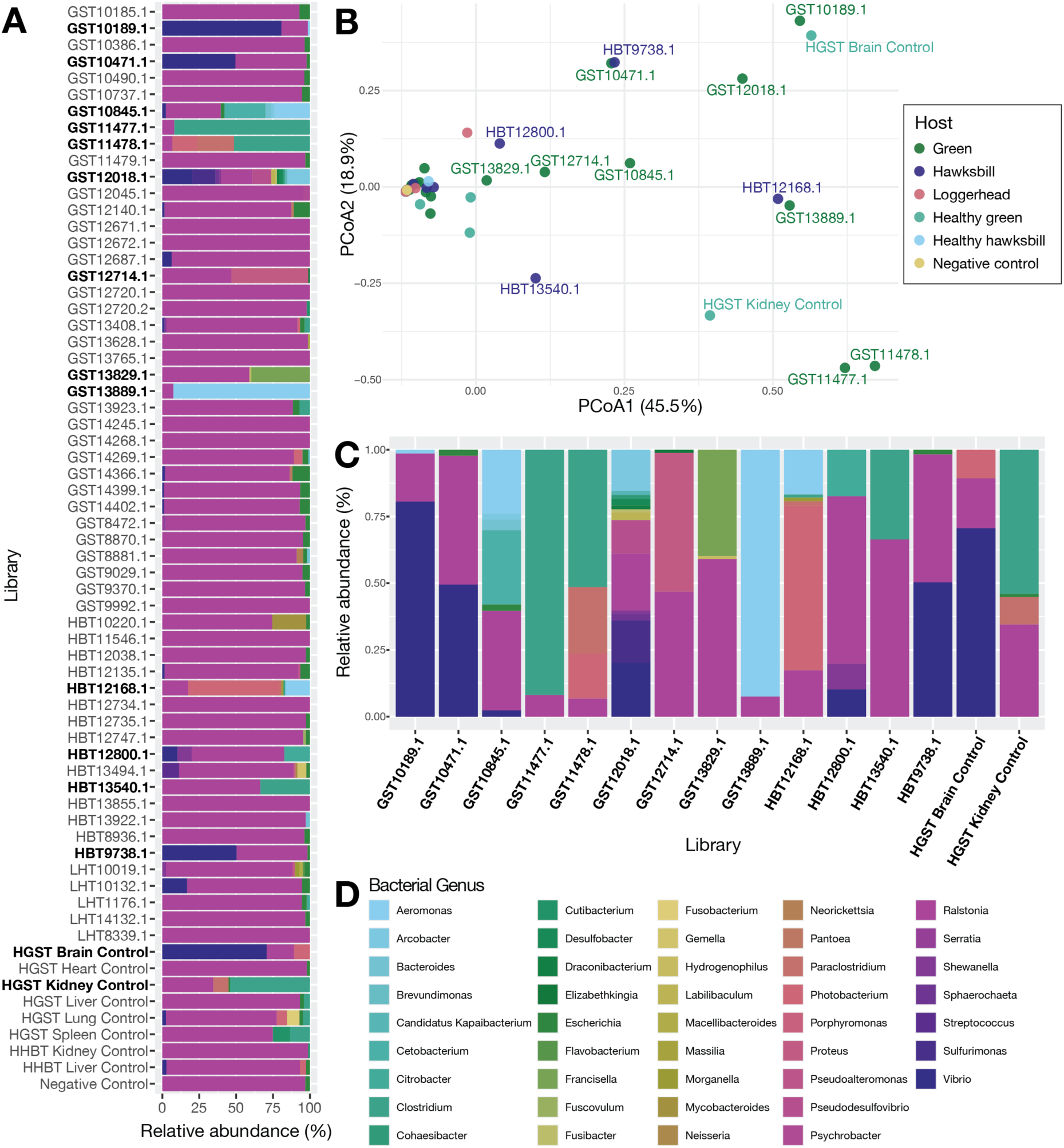
Marine turtle bacterial taxa profiles at the genus level. (A) Relative abundance plot of all libraries with distinct bacterial profiles in bold font. (B) Beta diversity measured by Bray-Curtis distance visualised using PCoA plot with distinct bacterial profile outlier libraries defined by MAD labelled. Key adjacent colours the sea turtle species. (C) Relative abundance plot of the 15 libraries with distinct bacterial profiles. (D) Colour key for bacterial genus for the relative abundance plots A and C.

A total of 15 cases were identified as having distinct (“outlier”) bacterial profiles that differed from the negative control (Figure 3B-C): GST10189.1, GST10471.1, GST10845.1, GST11477.1, GST11478.1, GST12018.1, GST12714.1, GST13829.1, GST13889.1, HBT12168.1, HBT12800.1, HBT13540.1, HBT9738.1, GST brain control, GST kidney control. Within these libraries, *Ralstonia* was present in lower relative abundance, and reads were instead dominated by those from the bacteria genera *Vibrio, Clostridium*, *Aeromonas*, *Proteus*, *Francisella*, *Photobacterium*, *Shewanella*, *Cetobacterium* and *Citrobacter* (Figure 3C). Four cases were dominated by *Vibrio* reads (GST10189.1, GST10471.1, HBT9738.1, GST brain control), another four by *Clostridium* reads (GST11477.1, GST11478.1, HBT13640.1, GST kidney control), while GST13889.1 was dominated by *Aeromonas* reads, GST12714.1 by *Proteus*, and GST13829.1 by *Franciscella*. The following libraries contained reads from multiple dominant bacterial genera: GST10845.1 with abundant *Cetobacterium* and *Aeromonas*, HBT12800.1 with *Citrobacter*, *Schwanella* and *Vibrio*, HBT12168.1 with *Aeromonas* and *Photobacterium*, and HBT13540.1 with *Vibrio* and *Schwanella* (Supplementary Table 4). Of note, the bacterial profile of GST12018.1 was more diverse than the others, including taxa such as *Arcobacter*, *Pseudodesulfovibrio*, *Ralstonia*, *Sulfurimonas* and *Vibrio*, consistent with marine turtle gastrointestinal-associated microbiota (Guo et al., 2022; Kuschke, 2022). However, because this library included gastric tissue, its profile likely reflects a gut-associated bacterial community rather than systemic infection.

Overall, we observed three distinct bacterial profiles in marine turtle metatranscriptomes: (i) the first likely reflects background environmental, reagent, or handling contamination as reflected by the abundance of *Ralstonia* and *Escherichia* (Salter et al., 2014; Xiao et al., 2019); (ii) the second reflects bacterial communities commonly observed in the natural microbiota of marine turtles (Kuschke, 2022); while (iii) the third was likely indicative of infection or colonisation of the host tissue either causing necrosis or decomposition (Dickson et al., 2011; Ebani, 2023). Indeed, this third group was also associated with pathology reports documenting pneumonia, sepsis, enteritis, coelomitis, tissue necrosis, severe blunt-force trauma to the carapace or tissue decomposition, with affected tissues included in the sequencing pool clearly impacting bacterial community composition (Supplementary Table 4).

Further investigation of the libraries with distinct bacterial profiles identified toxin-associated transcripts in five of the 15 cases (GST11477.1, GST11478.1, GST13889.1, HBT12168.1, HBT9738.1). The toxin transcripts identified included pore-forming or cytolytic toxins such as hemolysin, leukocidin, zona occludens toxin (Zot), exotoxin A, delta endotoxin, and toxin YoeB (Supplementary Table 5). While the presence of bacterial taxa cannot be directly linked to cause of disease, detection of toxin-associated transcripts suggests that some bacteria were transcriptionally active and expressing genes with potential roles in virulence or host damage.

### Viruses of marine turtles

A diverse set of viruses was observed in the marine turtle libraries. Because of their potential relevance to host pathology and disease we focused on likely vertebrate-associated viruses. The likely invertebrate-associated viruses detected, which will not be analysed further, came from the families *Botourmiaviridae*, *Mitoviridae*, *Partitiviridae*, *Phenuiviridae*, *Pseudototiviridae*, *Sobemoviridae*, *Tombusviridae*, *Totiviridae*, *Narnaviridae,* and *Fiersviridae*, as well as viruses assigned to the order *Formycovirales* (Supplementary Table 6). Although DNA viruses from the families *Adintoviridae* and *Herpesviridae* were also detected, these represented endogenous viral elements within in the host genome rather than active viral infection and hence were not analysed further.

### Chelonid hepaciviruses

The full genome of a novel hepacivirus, as well as sequences identical to the previously characterised hepacivirus sp. (GenBank accession UVD41733.1), were recovered from three marine turtle libraries: HBT12734.1, GST liver control and GST8472.1. The green turtle hepacivirus sequences identified here (GST8472.1 and GST liver control) exhibited 96.0% amino acid identity in their polyprotein to each other and to hepacivirus sp. The hawksbill turtle hepacivirus identified (HBT12734.1) had 76% amino acid identity to hepacivirus sp., thereby compatible with the host jumping of hepaciviruses among marine turtles.

Interestingly, these viruses formed a distinct marine turtle hepacivirus clade present in both the Australian Pacific Ocean and the South China Sea (Figure 4A). While individuals HBT12734.1 and GST8472.1 both had enteritis in their pathology reports, it is unlikely that marine turtle hepaciviruses are directly associated with disease. In particular, hepacivirus sp. was sampled from a healthy green turtle from Hainan, China, and the presence of the virus in the GST liver control suggests that it is an incidental finding rather than being directly disease associated. Interestingly, hepacivirus abundance was highest in the GST liver control (49.29TPM) followed by HBT12734.1 (3.48TPM) and GST8472.1 (2.20TPM).

**Figure 4.**
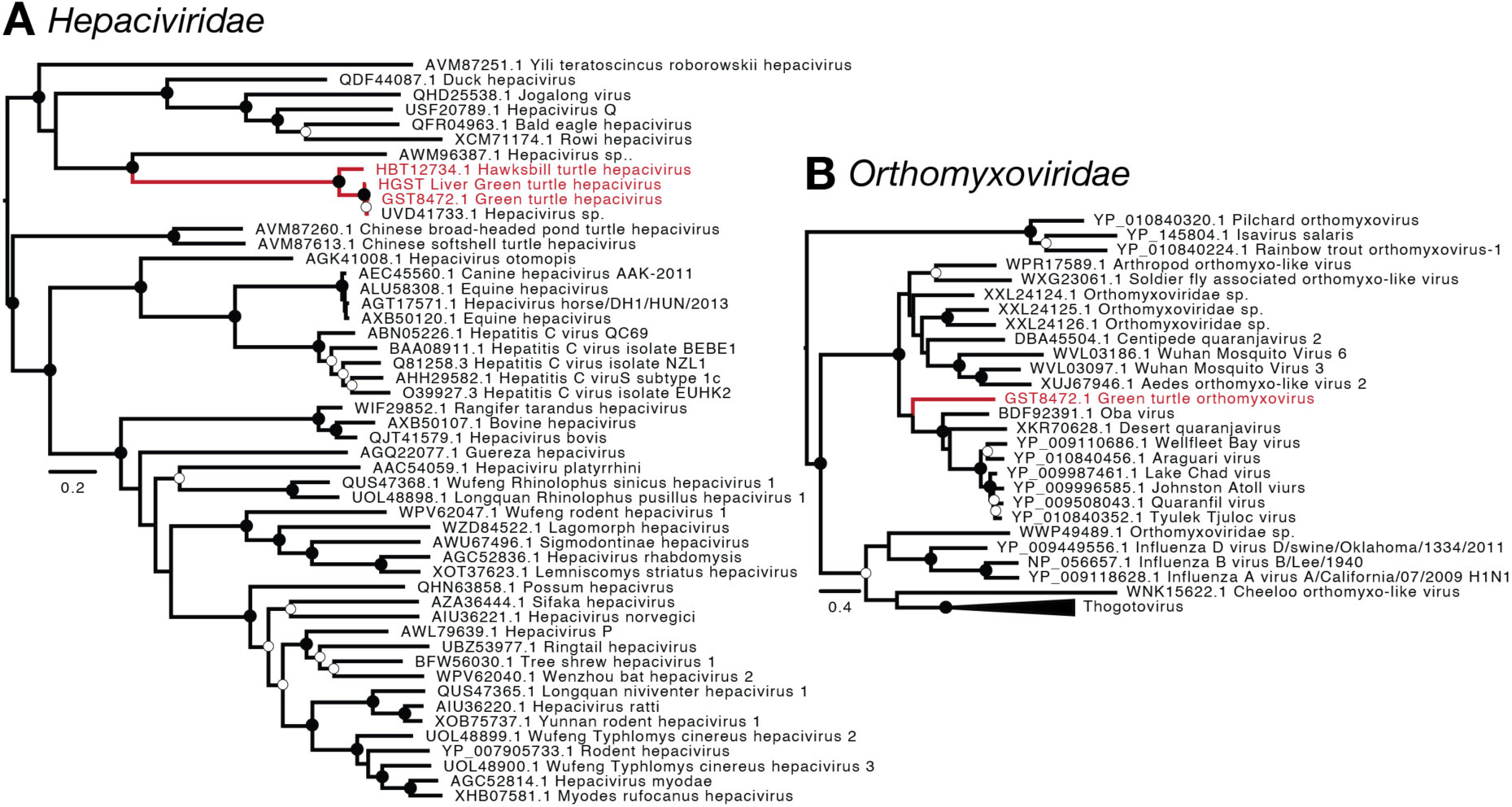
ML phylogenetic trees for the *Hepaciviridae* and *Orthomyxoviridae* based on the RdRp. (A) Novel hawksbill turtle hepacivirus from library HBT12734.1, and the previously identified green turtle Hepacivirus sp. (UVD41733.1) detected in the GST liver control and library GST8472.1. (B) Novel marine turtle orthomyxovirus found in GST8472.1. Viruses detected in this study are annotated in red. Mid-point rooted ML trees with nodes ≥95% bootstrap support annotated with black circles (●) and those 70-94% bootstrap support with white circles (○). Scale bar represents the estimated number of amino acid substitutions per site.

### Green turtle orthomyxovirus

Two partial polymerase basic protein 1 (PB1; encodes the orthomyxovirus RdRp) sequences and a polymerase acidic protein (PA) sequence were recovered from a green turtle (GST8472.1), with a combined abundance of 0.86TPM. Phylogenetic analysis of the PB1 gene indicated that green turtle orthomyxovirus is a member of the genus *Quaranjavirus* within the *Orthomyxoviridae* (Figure 4B). Although quaranjaviruses are thought to utilise ticks as vectors and have been identified in avian and human hosts, their host range and pathogenic potential remains poorly understood (Sameroff et al., 2021). Green turtle orthomyxovirus exhibited 47% amino acid identity with oba virus (BDF92391.1) detected in the blood-feeding mite *Dermanyssus hirundinis*. Oba virus was also the closest BLASTx match across the full PA gene, with 31% amino acid sequence identity. Phylogenetic analysis placed green turtle orthomyxovirus as the basal lineage in a clade containing oba virus and a group of closely related avian-associated quaranjaviruses, including Wellfleet Bay virus, Lake Chad virus and Johnston Atoll virus (Figure 4B). Notably, library GST8472.1 comprised brain, spleen, liver, kidney, lung, and plaque serosa. Although no parasites were detected in this library, it did contain sequences from green turtle hepacivirus. The primary cause of death reported for GST8472.1 was gastric torsion, such that these viruses most likely reflect coinfection than representing a direct cause of disease.

### Chelonid astrovirus

A novel marine turtle astrovirus was detected in both hawksbill and loggerhead turtles, with virus abundance ranging between 0.87-45.1TPM across positive libraries: HBT12168.1, 0.87TPM; LHT14132.1, 1.09TPM; HBT13540.1, 11.5TPM; LHT10019.1, 45.1TPM. Comparison of ORF1b RdRp sequences from these libraries revealed 98.5%-100% amino acid identity which suggests the detection of the same virus across all four libraries. A full astrovirus genome was recovered from HBT13540.1. The closest virus identified to the novel marine turtle astrovirus was the fish-associated Wuhan Japanese halfbeak astrovirus (AVM87484.1), yet with only 38% amino acid identity (Figure 5A). The broad presence of this marine turtle astrovirus in two species of turtles and across multiple individuals suggests that it likely represents a true turtle infection and not is associated with the fish component of turtle diet. In addition, libraries LHT14132.1 and LHT10019.1 exclusively comprised pooled lung and liver samples with no dietary tissue, yet both remained positive for the virus. In three of these cases, gut impaction suggested abnormal gastrointestinal function, and in the fourth case, anorexia and metabolic bone disease was reported (Supplementary Table 1). Astrovirus-positive libraries were associated with cases gastrointestinal upset, although further virus isolation and investigation is required to determine whether astrovirus infection contributed to these pathological findings or simply represents an incidental finding.

**Figure 5.**
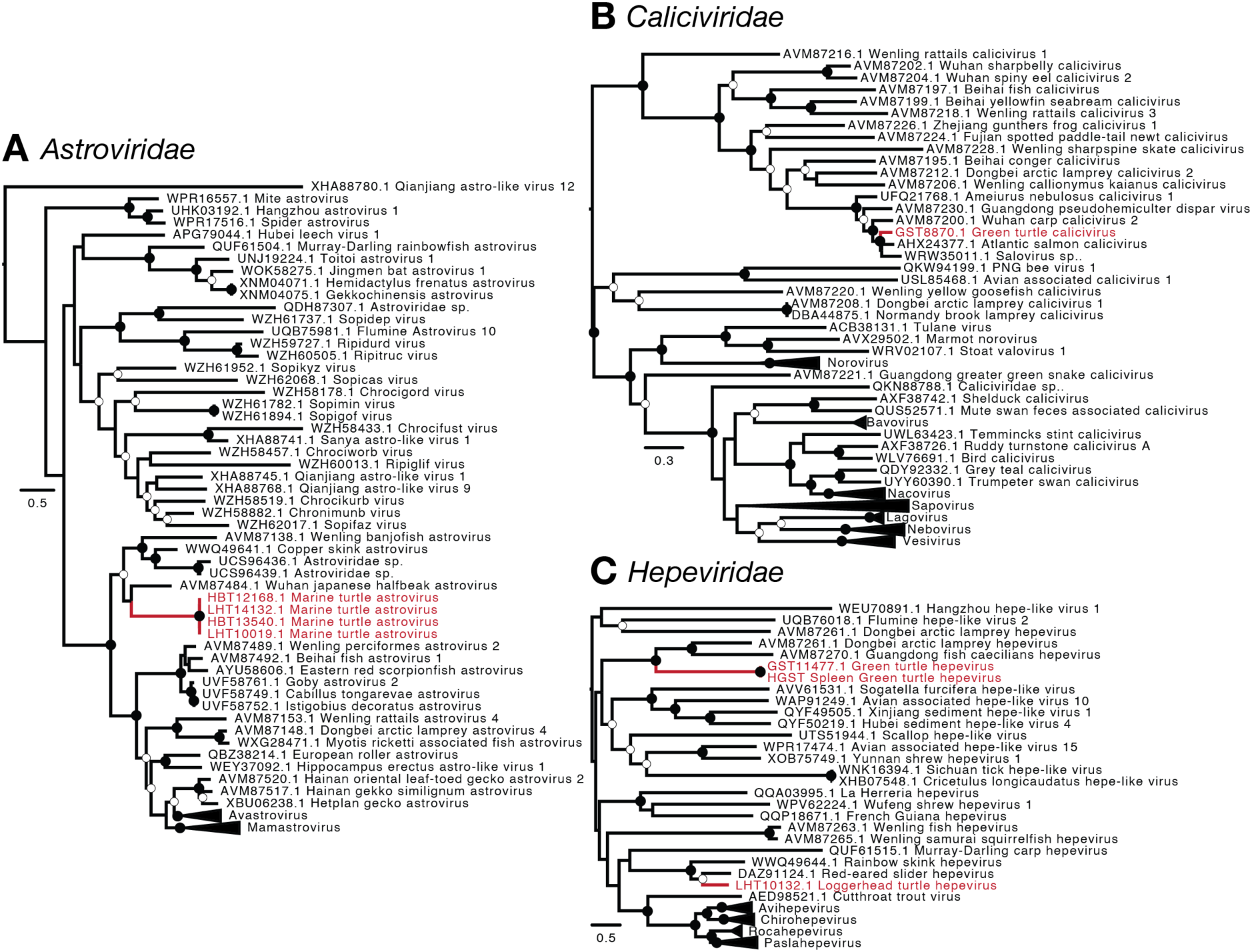
ML phylogenetic trees of the *Astroviridae*, *Caliciviridae* and *Hepeviridae* based on the RdRp. (A) The novel marine astrovirus was present in four marine turtle libraries: HBT12168.1, LHT14132.2, HBT13540.1, LHT10019. (B) Novel green turtle calicivirus found in library GST8870.1. (C) Novel marine turtle hepevirus from libraries GST11477.1, GST spleen control and LHT10132.1. Viruses detected in this study are annotated in red. Mid-point rooted ML trees with nodes ≥95% bootstrap support annotated with black circles (●) and those 70-94% bootstrap support with white circles (○). Scale bar represents the estimated number of amino acid substitutions per site.

### Green turtle calicivirus

A 1105bp partial RdRp of a novel green turtle calicivirus was detected at an abundance of 0.32TPM in library GST8870.1, sharing 83% RdRp amino acid identity with Atlantic salmon calicivirus (AHX24377.1) and 81% similarity with Wuhan carp calicivirus 2 (AVM87200.1). Although this novel calicivirus is found in a clade of fish associated viruses (Figure 5B), it is unlikely to represent dietary contamination as the sequencing library GST8870.1 comprises lung, liver and kidney samples with no dietary tissues and was also negative for fish reads in the Kraken2 analysis. The primary cause of death for the individual GST8870.1 was granulomatous pneumonia of fungal aetiology with microbial culture revealing *Metarhizium anisoplia*, a widely distributed soil-inhabiting fungus reported as a cause of granulomatous pneumonia in alligators. Interestingly, however, *M. anisoplia* was not detected using total RNA sequencing. We therefore interpret the green turtle calicivirus to be an incidental finding that is unlikely to have played a role in turtle mortality.

### Chelonid hepeviruses

The full genome of a novel green turtle hepevirus and a partial RdRp sequence of a novel loggerhead turtle hepevirus were also identified (Figure 5C). The newly identified loggerhead turtle hepevirus was detected in LHT10132.1 with a virus abundance of 1.41TPM. Identical green turtle hepevirus sequences were detected in two green turtle libraries, GST11477.1 and the GST spleen control, at abundances of 4.98TPM and 14.11TPM, respectively. These sequences were distantly related to other vertebrate-associated hepeviruses, including Dongbei arctic lamprey hepevirus (AVM87261.1; 27% amino acid identity). Interestingly, the loggerhead turtle hepevirus occupied a very different phylogenetic position, falling within a reptilian clade and exhibiting 65% amino acid identity to its closest relative, red-eared slider hepevirus (DAZ91124.1) (Figure 5C). The phylogenetic placement of the loggerhead turtle hepevirus within a reptile-associated lineage supports infection of the turtle host, but provides no evidence of disease association as none have been reported in the viruses within this clade. As the GST spleen control contained no dietary tissue or fish reads, both the green turtle and loggerhead turtle hepeviruses are neither diet associated nor linked to disease, and hence are more likely part of the natural marine turtle virome.

### Green turtle picobirnavirus

A partial 1515bp RdRp sequence of a green turtle picobirnavirus was found with an abundance of 0.2TPM in a green turtle (GST12018.1) that was reported to die of sepsis. No additional picobirnavirus-associated genome segments were found in this library. The green turtle picobirnavirus shared most amino acid similarity (47% RdRp amino acid identity) to picobirnavirus sp. (QXV86699.1) previously detected in rat faeces from China, and fell within a clade of mainly mammalian picobirnaviruses in phylogenetic analysis (Figure 6A). However, because only a partial RdRp sequence was recovered from a single library, its association with the turtle host and role in disease remains uncertain. In addition, members of the *Picobirnaviridae* infect a wide variety of hosts and environments, and also likely represent bacteriophage (Sadiq et al., 2024).

**Figure 6.**
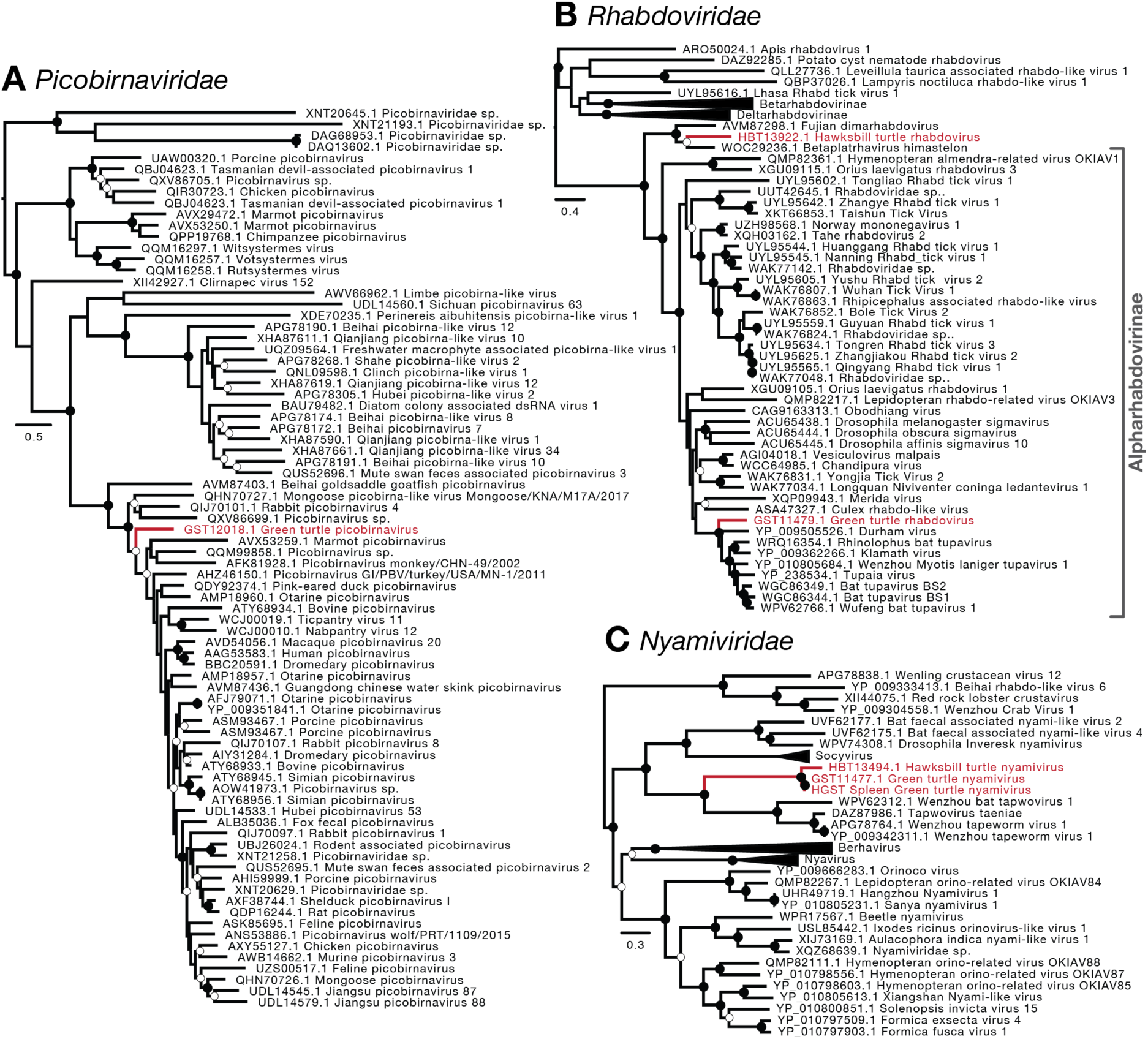
ML phylogenetic trees of the *Picobirnaviridae*, *Rhabdoviridae* and *Nyamiviridae* based on the RdRp. (A) Novel green turtle picobirnavirus from library GST12018.1. (B) Novel green turtle rhabdovirus from library GST11479.1. (C) Novel green turtle nyamiviruses present in three libraries: HBT13494.1, GST11477.1, healthy spleen green turtle. Viruses detected in this study annotated in red. Mid-point rooted ML trees with nodes ≥95% bootstrap support annotated with black circles (●) and those 70-94% bootstrap support with white circles (○). Scale bar represents the estimated number of amino acid substitutions per site.

### Chelonid rhabdoviruses

Two distinct turtle rhabdoviruses were identified here, from a hawksbill turtle (HBT13922.1) and a green turtle (GST11479.1), with virus abundances of 1.93TPM and 61.86TPM, respectively (Figure 6B). The hawksbill turtle rhabdovirus was most closely related to Himastelon rhabdovirus 1 (WOC29236.1) isolated from the parasitic flatworm *Himasthla elongata* (27% RdRp amino acid identity), and to Fujian dimarhabdovirus (AVM87298.1) found in a spotted paddle-tail newt from China (35% RdRp amino acid identity) (Figure 6B). While its similarity to the Fujian dimarhabdovirus indicates infection of the hawksbill turtle host, the detection of hapalotrema reads in HBT13922.1 and its relation to Himastelon rhabdovirus 1 means that a virus-parasite association cannot be excluded. In contrast, the green turtle rhabdovirus fell within the subfamily *Alpharhabdovirinae*, previously reported in both vertebrate and invertebrate hosts, and was most similar (51% RdRp amino acid identity) to Durham virus (YP_009505526.1), a tupavirus associated with bird infection (Figure 6B). As GST11479.1 contained no detected parasite reads it is likely that this virus infects the turtle host. The primary causes of death reported for HBT13922.1 and GST11479.1 were oesophagitis and entanglement, respectively, suggesting no clear association with virus pathology.

### Cheloniid nyamiviruses

Two novel nyamiviruses were identified this study, falling adjacent in a phylogenetic analysis to members of the genus *Tapowovirus* (*Nyamiviridae*) that are commonly reported in tapeworms (Figure 6C). Partial RdRp fragments were recovered from libraries GST11477.1 (abundance of 0.49T), HBT13494.1 (abundance of 0.41TPM) and the GST spleen control (abundance of 2.84TPM), with the longest sequences being 8157bp, 1730bp and 1627bp, respectively. Based on the nucleotide alignment of the RdRp, the partial 1627bp sequence identified in the healthy green turtle spleen control was identical to the sequence from GST11477.1, such that they likely represent the same virus. The virus with the greatest amino acid similarity to these marine turtle nyamiviruses was Wenzhou bat tapwovirus 1 (WPV62312.1) isolated from bat faeces (to which other tapeworm-associated viruses were closely related), sharing 49% amino acid identity. It is important to note that because all three nyamivirus-positive libraries are also parasite-positive, it is unclear whether this virus infects the marine turtle host or its associated parasites.

### Microbial co-detection within the pathobiome

A notable feature of our analysis was the co-detection of multiple infectious agents across the pathobiome in 21 sequencing libraries (Figure 7A), with 15 showing evidence of inter-kingdom co-infections involving viral, bacterial and eukaryotic parasite taxa (Figure 7B). The frequent identification of multiple candidate infectious agents complicates the attribution of disease to a single agent when only considering the metagenomic data alone. Virus diversity varied by host species, with a diverse range of viruses from seven virus families detected in the green turtle, contrasting with only two virus families identified in the hawksbill turtle and two in the loggerhead turtle (Figure 7C). However, as green turtles represent the largest host group in the data set (n=54), this likely reflects sampling bias rather than increased host species susceptibility.

**Figure 7.**
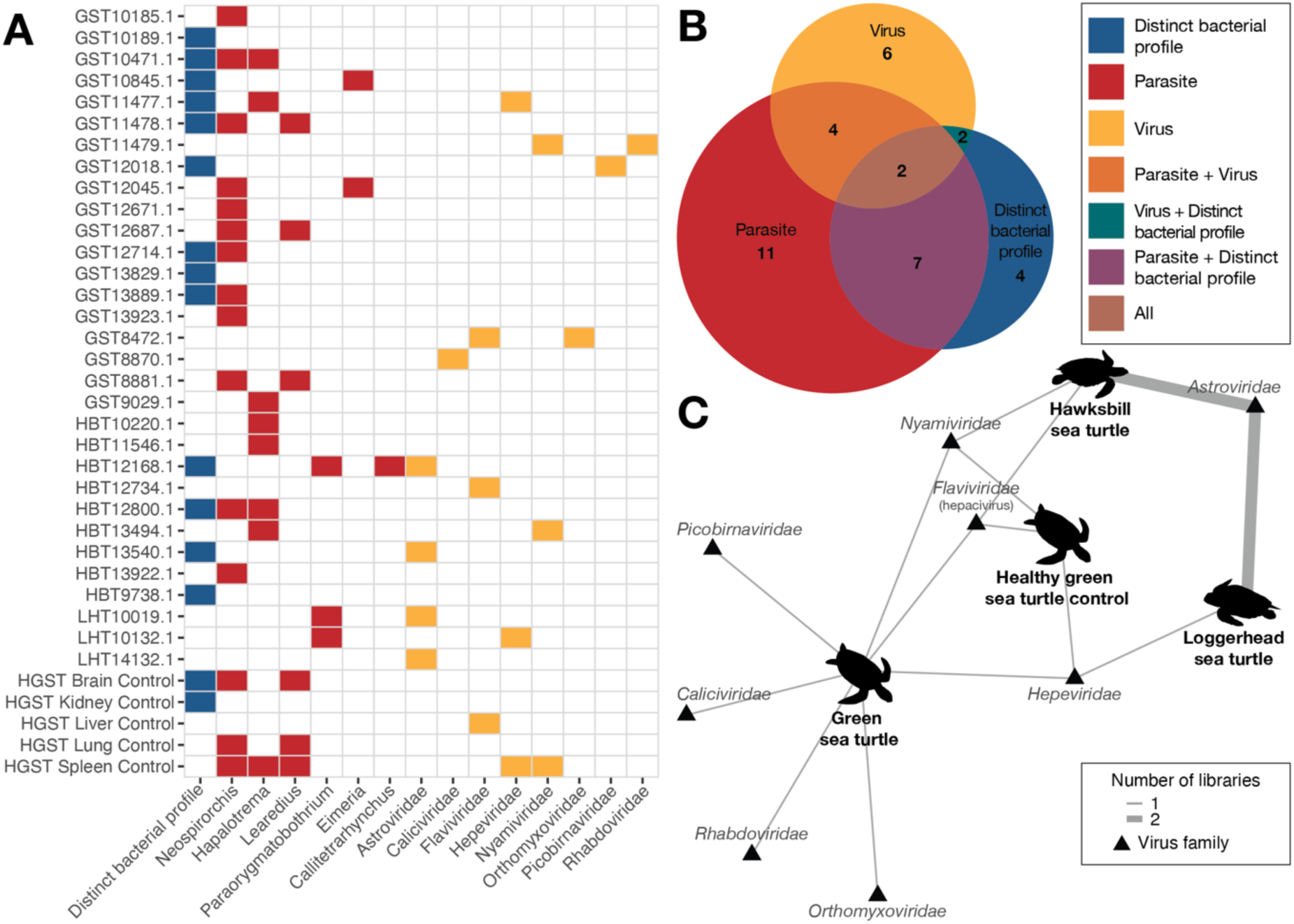
Summary of marine turtle infectious agents in this study. (A) Heatmap of sequencing libraries with reported infectious agents. (B) Euler diagram displaying the number of libraries with eukaryotic parasites, distinct bacterial profiles and virus infection, and the prevalence of their co-infections coloured as follows: distinct bacterial profile (blue), parasite (red), virus (yellow), parasite and virus coinfection (orange), virus and distinct bacterial profile (green), parasite and distinct bacterial profile (purple), all (brown). (C) Virus network displaying three species of marine turtles sampled in this study and the healthy green sea turtle control libraries.

## Discussion

Marine turtles are a global keystone species that play a critical role in maintaining the ecological balance of sea grass beds and coral reefs across the world. Herein, we show that marine turtles harbour a great diversity of microbes and parasitic life including eukaryotic parasites, bacteria and viruses. While many of these agents are likely to represent incidental findings, some may also possess the potential to cause disease.

Across the marine turtle pathobiome data set, the most prominent metatranscriptomic signal was from eukaryotic parasites, particularly *Neospirorchis* and *Hapalotrema*, which have been previously reported across all marine turtle species (Chapman et al., 2019). Consistent with previous studies, *Neospirorchis* and *Hapalotrema* are common in marine turtles, perhaps persisting at a low burden, but contributing to disease when parasite loads are high or when egg deposition causes tissue damage.

Also of note was that a subset of marine turtle metatranscriptomic libraries contained bacterial communities that were markedly distinct from the *Ralstonia*-dominated background profile observed across most libraries. Several of these distinct libraries were dominated by opportunistic bacterial genera previously associated with marine turtle infections and were detected in cases with pathology consistent with bacterial infection. However, because contamination, tissue decomposition, necrosis, and carapace trauma may collectively influence bacterial abundance and composition, these findings should be interpreted with caution. Indeed, rather than confirming bacterial-associated pathology, this data illustrates that total RNA sequencing can detect differences in bacterial communities within a host that may warrant further investigation.

Twelve novel marine turtle-associated viruses were identified across eight virus families. Of particular note was that, within their respective families, the marine turtle viruses either formed distinct lineages characterised by relatively long branch lengths (i.e., relatively low amino acid similarities to their closest relatives) or clustered adjacent to other marine vertebrates, particularly fish, indicative of relatively long periods of independent evolution. Interestingly, we also observed both cross-species transmission, wherein a virus was detected in multiple host species, and instances of distinct virus lineages from the same virus family associated with different cheloniid species. For example, the novel marine turtle astrovirus was detected in both hawksbill and loggerhead turtles across four libraries, indicative of relatively recent host-jumping. In contrast, phylogenetically distinct rhabdoviruses (∼15% RdRp amino acid sequence similarity) were identified in the green turtle and hawksbill turtle, while distinct hepeviruses (30% amino acid identity) were identified in the green turtle and loggerhead turtle, with the latter most closely related to other reptilian hepeviruses. While such distinct viruses may represent host species specificity for these viruses, it could similarly reflected limited sampling, the uneven representation of turtle species, or low virus prevalence within our sample population. Determining whether these marine turtle viruses (in the *Orthomyxoviridae*, *Caliciviridae*, *Hepeviridae*, and *Picobirnaviridae*) only identified in single host species are capable of cross-species transmission would require further experimentation such as virus isolation, culture and transfection assays which are beyond the scope of this study.

Importantly, the marine turtle viruses characterised in this study seem to be commensal in nature and not clearly associated with disease. Indeed, none of the pathology reports attribute virus infection as the primary cause of turtle mortality in this study. Further study on co-infection and disease dynamics is necessary to evaluate the true impact of these viruses on turtle health, including whether they are endemic to marine turtles and proliferate better in immunocompromised hosts such as those in this study brought in due to injury, infection or other illness.

In sum, our study highlights the complexity of the marine turtle pathobiome and suggests that disease in these hosts is unlikely to be explained by a single infectious agent. Rather, our data are compatible with parasite burden, proliferation of opportunistic bacterial communities, and diverse viral infections interacting with host condition, injury, immune status, and environmental stressors to shape disease outcomes. Notably, co-infection across the marine turtle pathobiome was commonplace within our data, highlighting the need to consider the synergistic effect of multiple infectious agents when investigating turtle health.

## Data Availability

The raw sequence reads generated here are available on the NCBI SRA under BioProject PRJNA1534595 and accession numbers SRR40894471-SRR40894407. All viral genomes and corresponding sequences assembled in this study have been deposited in GenBank under accession numbers XXXX-YYYY.

## Acknowledgements

The Australian Registry of Wildlife Health (the Registry) is a 40-year science program of Taronga Conservation Society, which contributes to the surveillance of marine species health in partnership with the New South Wales Departments of Primary Industries and Regional Development and Climate Change, Energy, the Environment and Water. We thank the previous and ongoing efforts of staff and volunteers in assisting with managing samples and data related to wildlife health programs, particularly Drs. Peter Johnson and Bryn Lynar, and Nicole Dobson, Allison Rubie, and Karina Hammond. We also recognize the University of Sydney Veterinary Pathology Diagnostic Services for their assistance in processing histopathology slides and the technical assistance provided by the Sydney Informatics Hub, a Core Research Facility of the University of Sydney.

## Supplementary Information

**Supplementary Figure 1.** Chelonid pathobiome composition based on relative abundance of classified Kraken reads.

**Supplementary Figure 2.** Eukaryotic parasite reads mapped against reference sequences. Blue graphs representing parasites with high read counts and mapping coverage.

**Supplementary Table 1.** Case overview of marine turtles including species, sex, age, date of death, location, and summary of pathology.

**Supplementary Table 2.** Data summary of sequencing libraries, pooling schedule, and sample tissue types.

**Supplementary Table 3.** Summary of parasite detections and read counts.

**Supplementary Table 4.** Overview of libraries with distinct bacterial profiles.

**Supplementary Table 5.** Bacterial toxin hits to the Toxinome database (Danov et al., 2023) and their read abundance.

**Supplementary Table 6.** Overview of virus contigs detected in this study.

