## Supplementary Figures S1-S2 for "Metatranscriptomic Identification of Co- and Incidental Infections in Marine Turtles Reveals Complex Disease Aetiologies"

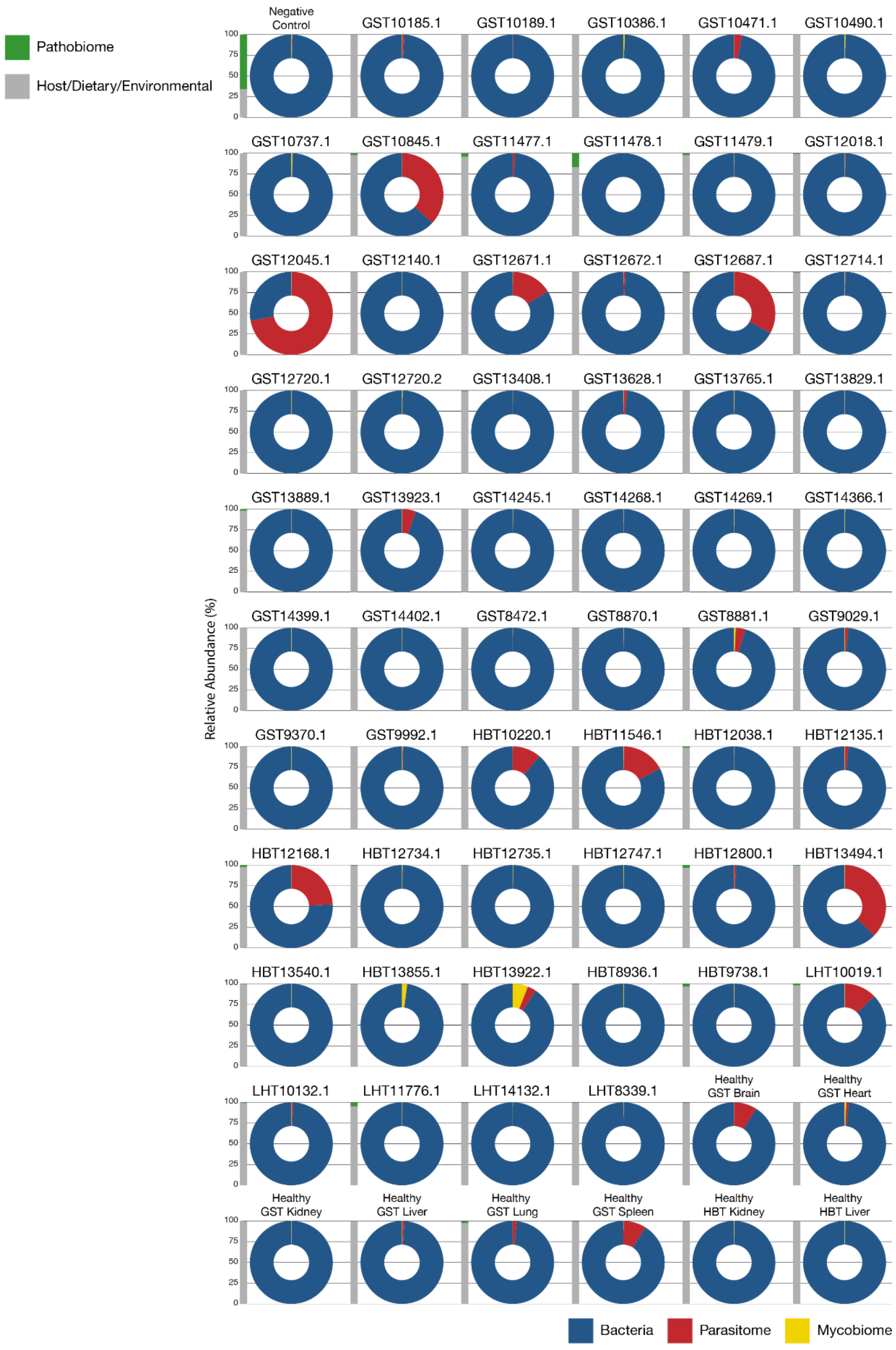


**Supplementary Figure 1.** Chelonid pathobiome composition based on relative abundance of classified Kraken reads.

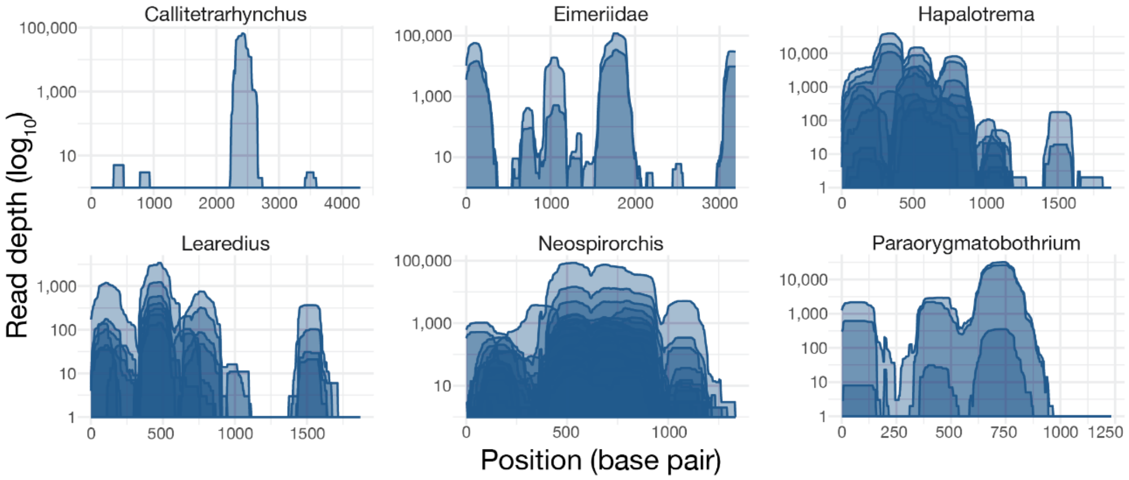


**Supplementary Figure 2.** Eukaryotic parasite reads mapped against reference sequences. Blue graphs representing parasites with high read counts and mapping coverage.
